# A nonsense mutation in the *COL7A1* gene causes epidermolysis bullosa in Vorderwald cattle

**DOI:** 10.1101/062968

**Authors:** Hubert Pausch, Simon Ammermüller, Christine Wurmser, Henning Hamann, Jens Tetens, Cord Drögemüller, Ruedi Fries

## Abstract

**Background:** The widespread use of individual sires for artificial insemination promotes the propagation of recessive conditions. Inadvertent matings between unnoticed carriers of deleterious alleles may result in the manifestation of fatal phenotypes in their progeny. Breeding consultants and farmers reported on Vorderwald calves with a congenital skin disease. The clinical findings in affected calves were compatible with epidermolysis bullosa.

**Results:** Pedigree analysis indicated autosomal recessive inheritance of epidermolysis bullosa in Vorderwald cattle. We genotyped two diseased and 41 healthy animals at 41,436 single nucleotide polymorphisms and performed whole-genome haplotype-based association testing, which allowed us to map the locus responsible for the skin disease to the distal end of bovine chromosome 22 (P=8.0×10^−14^). The analysis of whole-genome re-sequencing data of one diseased calf, three obligate mutation carriers and 1682 healthy animals from various bovine breeds revealed a nonsense mutation (rs876174537, p.Arg1588X) in the *COL7A1* gene that segregates with the disease. The same mutation was previously detected in three calves with dystrophic epidermolysis bullosa from the Rotes Höhenvieh cattle breed. We show that diseased animals from Vorderwald and Rotes Höhenvieh cattle are identical by descent for an 8.72 Mb haplotype encompassing rs876174537 indicating they inherited the deleterious allele from a recent common ancestor.

**Conclusions:** Autosomal recessive epidermolysis bullosa in Vorderwald and Rotes Höhenvieh cattle is caused by a nonsense mutation in the *COL7A1* gene. Our findings demonstrate that recessive deleterious alleles may segregate across cattle populations without apparent admixture. The identification of the causal mutation now enables the reliable detection of carriers of the defective allele. Genome-based mating strategies can avoid inadvertent matings of carrier animals thereby preventing the birth of homozygous calves that suffer from a painful skin disease.

## Background

Vorderwald cattle are mainly kept in the southwestern part of Germany. The breeding population consists of less than 10,000 cows of which 50% are artificially inseminated [1]. In spite of the small number of breeding animals, the effective population size (N_e_) of Vorderwald cattle is relatively high (N_e_=130) [1]. Extensive cross-breeding of Vorderwald cows with sires from the Ayrshire, Red Holstein and Montbéliarde breeds was performed in the past 50 years to improve milk, beef and conformation traits [2, 3, 4]. The contributions of foreign ancestors to the genomic composition of the current Vorderwald cattle population are in excess of 60% [3].

The availability of genome-wide genotyping and sequencing data facilitates the mapping of Mendelian traits [5, 6, 7]. Most genetic disorders in cattle populations follow an autosomal recessive inheritance likely because the widespread use of individual bulls for artificial insemination promotes the propagation of recessive alleles [8, 9, 10]. Inadvertent matings between unnoticed carriers of harmful recessive alleles may occur frequently particularly in small populations and can result in the manifestation of fatal phenotypes in their progeny [11].

Inherited disorders had been rarely reported in Vorderwald cattle. The cross-breeding of Vorderwald cows with sires from foreign breeds possibly prevented recessive deleterious alleles from occurring in the homozygous state. However, preferred selection of animals carrying foreign haplotypes with favourable effects on breeding objectives (*e.g.*, [4, 12]) may cause local inbreeding affecting specific regions of the genome and can result in the phenotypic manifestation of recessive alleles that were absent in the unadmixed population [13, 14, 15, 16].

Epidermolysis bullosa (EB) is an inherited disorder of the connective tissue that had been observed in many species including cattle [17]. Affected individuals suffer from skin blisters and high skin fragility because of an impaired adherence of the epidermis to the underlying dermis [17]. There is no cure for EB. However, skin and wound care may alleviate the clinical symptoms [18]. Depending on the severity of the clinical manifestations, EB may be fatal. Causal mutations for bovine EB had been identified in the *COL7A1* (OMIA 000341-9913) [19], *ITGB4* (OMIA 001948-9913) [20, 21], *KRT5* (OMIA 000340-9913) [22], *LAMA3* (OMIA 001677-9913) [23] and *LAMC2* (OMIA 001678-9913) [24] genes.

Herein we present the phenotypic and molecular-genetic characterisation of autosomal recessive epidermolysis bullosa in Vorderwald cattle. We exploit whole-genome genotyping and sequencing data to identify the causal mutation for the skin disease in the *COL7A1* gene. We show that a haplotype encompassing the deleterious allele, unexpectedly, also segregates in Rotes Höhenvieh cattle.

## Results

### Phenotypic manifestation and inheritance of a painful skin disease in Vorderwald cattle

Twenty-five Vorderwald calves (17 male, 6 female, 2 of unknown sex) with congenital skin lesions were born between the years 2004 and 2015. The calves had extensive hairless, reddened and bloody regions at the fetlocks (Figure 1a-c). Although the calves were able to stand properly, they were reluctant to move likely due to painful lesions at the fetlocks. Cartilage deformities, hairless and bloody sites were observed at the calves’ ears (Figure 1b). Based on the phenotypical appearance of the calves, the tentative diagnosis of epidermolysis bullosa (EB) was made. Since pathological and histological data were not available, a more precise classification of the disease was not possible [25]. All calves were euthanized shortly after birth by accredited veterinarians because of poor prognosis and with no prospect of improvement [17]. Breeding consultants collected ear tissue samples for genetic analyses from two affected calves.

**Figure 1.**
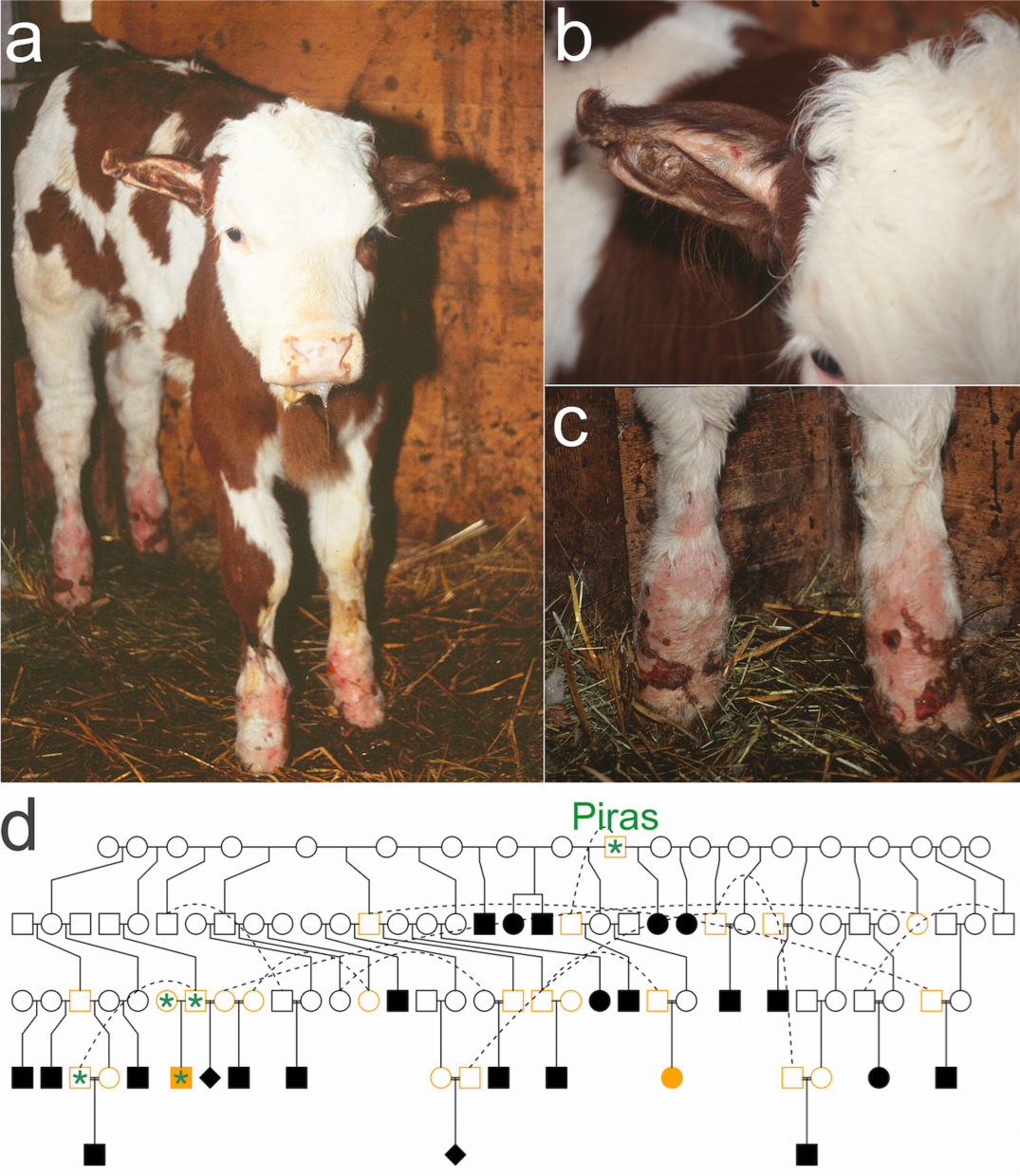
A Vorderwald calf with a painful skin disease. A few days old calf with reddened and bloody sites at its fetlocks and ear cartilage aberrations (a-c). The relationship between the Vorderwald x Montbéliarde crossbred bull Piras and 25 calves with epidermolysis bullosa (d). Circles, rectangles and diamonds represent females, males and animals of unknown sex, respectively. Filled symbols denote diseased animals. Orange and green colours denote animals that were available for genotyping and sequencing.

In order to determine the mode of inheritance of EB, we inspected pedigree records of 25 affected calves. The Vorderwald x Montbéliarde crossbred bull Piras (born in 1996) was in the paternal path of all and in the maternal path of 15 affected calves, suggesting recessive inheritance (Figure 1d). Six diseased calves were sired by Piras. However, Piras was absent in the maternal path of ten affected calves possibly due to incomplete pedigree records. It is also possible that the mutation occurred several generations before Piras. Piras inherited 50%, 10% and 1% of its genes from ancestors of the Montbéliarde, Holstein and Ayrshire breed, respectively, suggesting that the EB-associated mutation may had been introgressed into Vorderwald cattle from a foreign breed. Forty-six ancestors of Piras including a Holstein bull (Max, born in 1975) and an Ayrshire bull (Sypland Officer, born in 1951) were in the maternal path of all affected calves.

### Epidermolysis bullosa maps to chromosome 22

In order to map the genomic region associated with EB, two affected calves, 14 parents of another 23 affected calves including Piras and 27 healthy artificial insemination bulls from the Vorderwald cattle breed were genotyped using medium-density genotyping arrays. Following quality control, genotypes of 41,436 SNPs were phased to obtain haplotypes for association testing. We assumed recessive inheritance and used the expected number of copies of the EB-associated haplotype (2, 1 and 0 for affected calves, parents of affected calves and control animals, respectively) as response variable in a linear regression model. To account for the relationship among the genotyped animals, we considered ten principal components in the regression analysis. A moderate inflation factor of 1.24 indicated that this corrective measure was mostly successful. Fifty-five haplotypes located at the distal end of BTA22 were significantly associated with EB (P<8.6×10^−7^, Figure 2a-b). There were no significantly associated haplotypes detected on chromosomes other than BTA22. The top association signals (P=8.0×10^−14^) resulted from 19 adjacent haplotypes located between 42,994,215 bp and 49,273,889 bp. The two diseased calves had a common 21.26 Mb segment of extended homozygosity at the distal end of BTA22 (between 40,113,694 bp and 61,378,199 bp) (Figure 2c) that was not found in the homozygous state in 41 unaffected animals, corroborating a recessive mode of inheritance.

**Figure 2.**
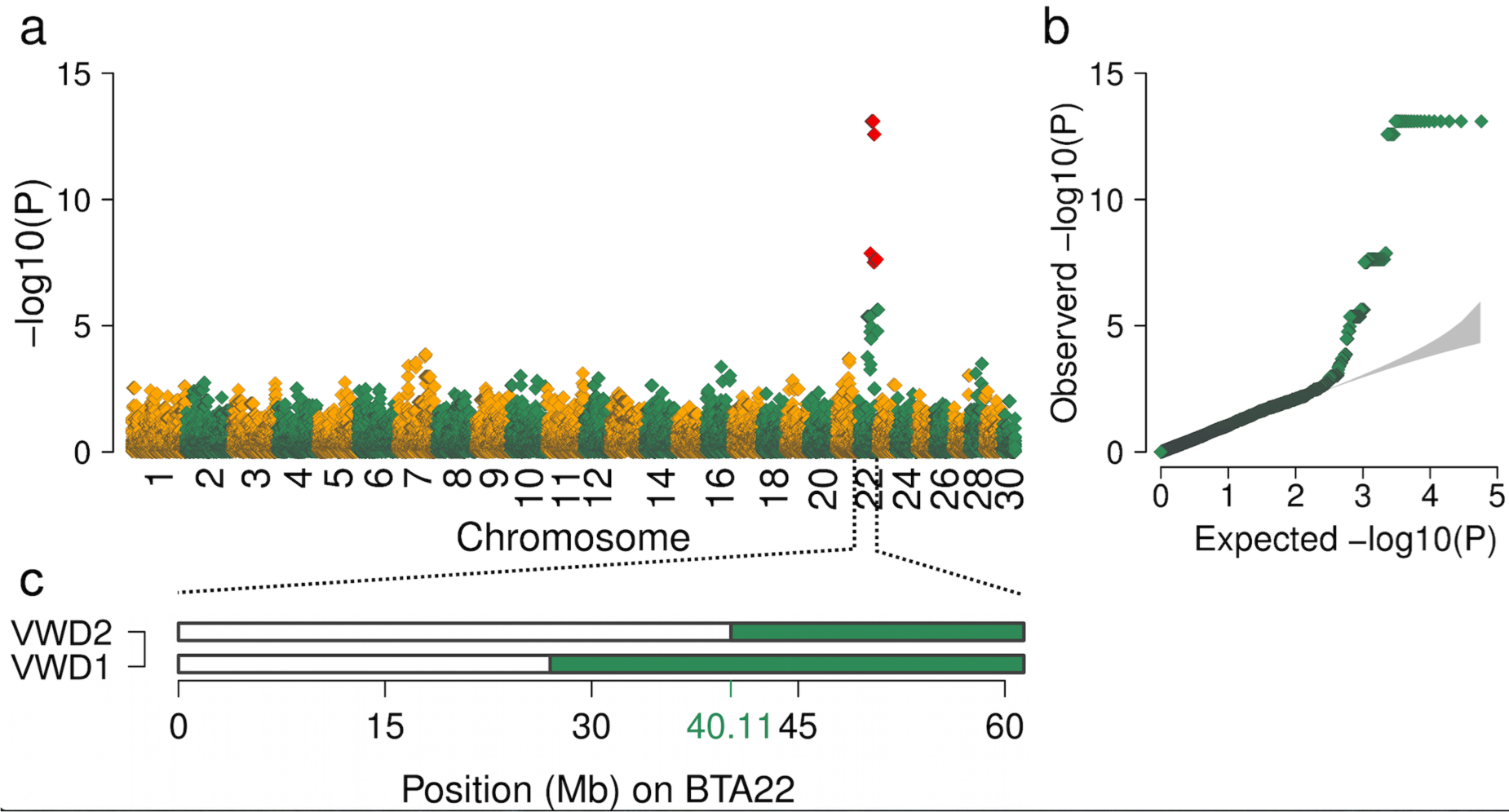
Haplotype-based association mapping of epidermolysis bullosa. Association of 57,837 haplotypes with epidermolysis bullosa (a). Red colour represents significantly associated haplotypes (P<8.64 x 10^−7^). Quantile-quantile plot (b). The grey shaded area represents the 95%-concentration band under the null hypothesis of no association. Homozygosity mapping in two affected Vorderwald (VWD) calves on bovine chromosome 22 (c). The green colour indicates segments of extended homozygosity in two calves with EB.

### A nonsense mutation in the *COL7A1* segregates with the skin disease

To identify the causal mutation for EB, an affected calf, its parents and the bull Piras were sequenced to an average read depth of 22.78x, 19.02x, 19.57x and 10.24x, respectively. To facilitate the identification of plausible candidate causal variants, we exploited sequence data of another 1682 animals from various bovine breeds that had been generated previously including three key ancestors from the Vorderwald cattle population. None of them was affected by EB. Multi-sample variant calling within the 21.26 Mb segment of extended homozygosity yielded genotypes at 71,169 single nucleotide and short insertion and deletion polymorphisms in four Vorderwald cattle. Of these, 8100 variants were compatible with recessive inheritance that is homozygous for the non-reference allele in the calf with EB and heterozygous in its parents and the bull Piras. However, 7398 of the compatible variants are most likely not causal for EB because they occurred in the homozygous state at least once in 1682 healthy animals. In conclusion, 702 variants were homozygous for the non-reference allele in an affected calf, heterozygous in obligate mutation carriers and not homozygous for the non-reference allele in healthy animals. Twenty candidate causal variants were splice site mutations or resided in protein coding regions of the genome (see Additional file 1), among these, a nonsense mutation in the *COL7A1* gene (*collagen type VII alpha 1*, Chr22:51873390, rs876174537, ENSBTAT00000025418:c.4762C>T, ENSBTAP00000025418:p.Arg1588X) that was previously detected in three Rotes Höhenvieh calves with dystrophic EB [19]. Sanger sequencing confirmed that both diseased Vorderwald calves were homozygous for the rs876174537 T-allele indicating that the same mutation causes EB in two distinct cattle breeds. The rs876174537 T-allele was not detected in 1682 sequenced animals from various bovine breeds other than Vorderwald.

We genotyped 86 healthy Vorderwald cattle at rs876174537 including 35 that had been genotyped with Illumina’s Bovine SNP50 genotyping array (Table 1). The rs876174537 polymorphism was in complete linkage disequilibrium with the EB-associated haplotype; 14 haplotype carriers were heterozygous at rs876174537 and twenty-one animals that did not carry the EB-associated haplotype, were homozygous for the reference allele. Of 79 artificial insemination bulls that were born between 1982 and 2014, eight were heterozygous mutation carries including seven sires of calves with EB, corresponding to a frequency of the deleterious allele of 5%. All artificial insemination bulls carrying the rs876174537 T-allele were descendants (sons or grandsons) of Piras.

**Table 1:** **Genotypes of 88 Vorderwald cattle** at the rs876174537 polymorphism **in** ***COL7A1***

| Animal group | N | Genotypes at the *COL7A1* variant |  |  |
| --- | --- | --- | --- | --- |
|  |  | CC | CT | TT |
| Calves with epidermolysis bullosa | 2 | - | - | 2 |
| Obligate carriers |  |  |  |  |
| - Dams of calves with EB | 7 | - | 7 | - |
| - Sires of calves with EB | 7 | - | 7 | - |
| Healthy bulls used in artificial insemination |  |  |  |  |
| - Carriers of the EB-associated haplotype | 0 | - | - | - |
| - Non-carriers | 21 | 21 | - | - |
| - Haplotype state unknown | 51 | 50 | 1 | - |

### A common haplotype encompassing the rs876174537 T-allele segregates in Vorderwald and Rotes Höhenvieh cattle

Menoud et al. [19] detected the rs876174537 T-allele in three Rotes Höhenvieh calves with recessive dystrophic EB. Two affected Vorderwald calves had an 8.72 Mb segment of extended autozygosity encompassing the rs876174537 T-allele in common with the diseased calves from Rotes Höhenvieh cattle indicating they inherited the EB-associated haplotype from a common ancestor (Figure 3). However, they had no common ancestors in their pedigrees likely because of incomplete pedigree records particularly for the diseased calves from Rotes Höhenvieh cattle [19, 26].

**Figure 3.**
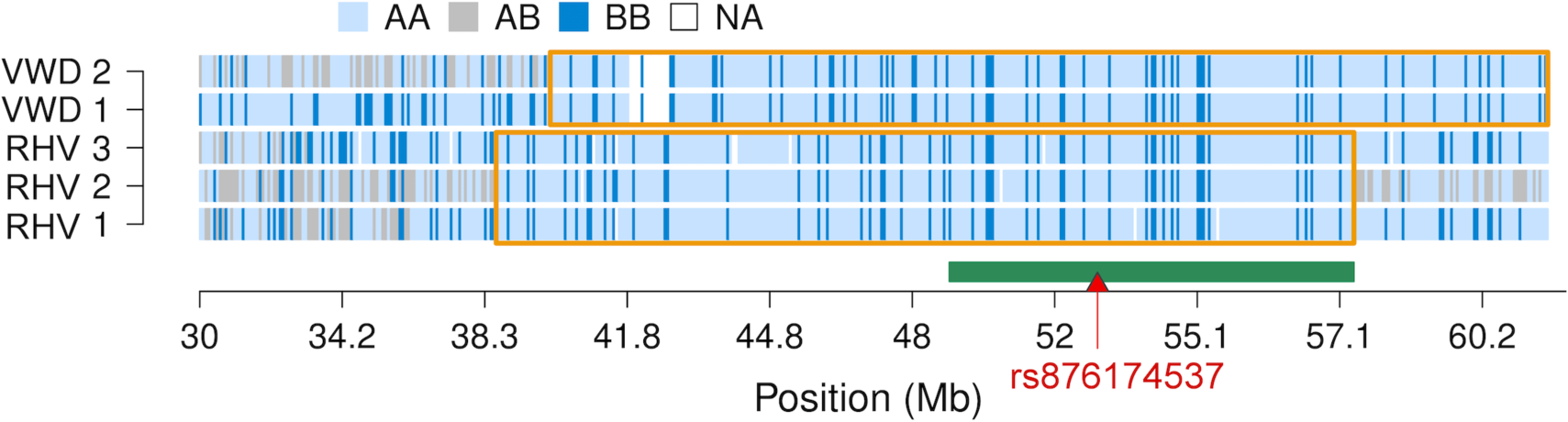
Homozygosity mapping on bovine chromosome 22 in five calves with epidermolysis bullosa. Light blue, grey and dark blue, respectively, represent homozygous, heterozygous and alternate homozygous genotypes in five calves with epidermolysis from Vorderwald (VWD) and Rotes Höhenvieh (RHV) cattle. The orange frames indicate common segments of extended homozygosity within breeds. The green bar represents a 8.72 Mb segment of extended homozygosity that was common in the diseased calves from both breeds. The red triangle represents the position of the rs876174537 T-allele (red triangle) in the *COL7A1* gene.

In an attempt to determine the origin of the EB-associated haplotype, we analysed array-derived genotypes of 714 animals from seven breeds including 43 Vorderwald and 12 Rotes Höhenvieh cattle (Figure 4). The top two principal components of the genomic relationship matrix separated the animals by breed. A high genetic heterogeneity was observed in Red Dairy Cattle which corroborates previous findings [27] (Figure 4a). Vorderwald and Rotes Höhenvieh cattle clustered next to each other. Moreover, Vorderwald cattle clustered close to Montbéliarde cattle likely because of the frequent cross-breeding in the past. An admixture of Vorderwald with Red Dairy and Holstein cattle was not apparent. The estimation of ancestry proportions using the *fastSTRUCTURE* software tool corroborated that Vorderwald, Rotes Höhenvieh and Montbéliarde cattle are closely related (Figure 4b).

**Figure 4.**
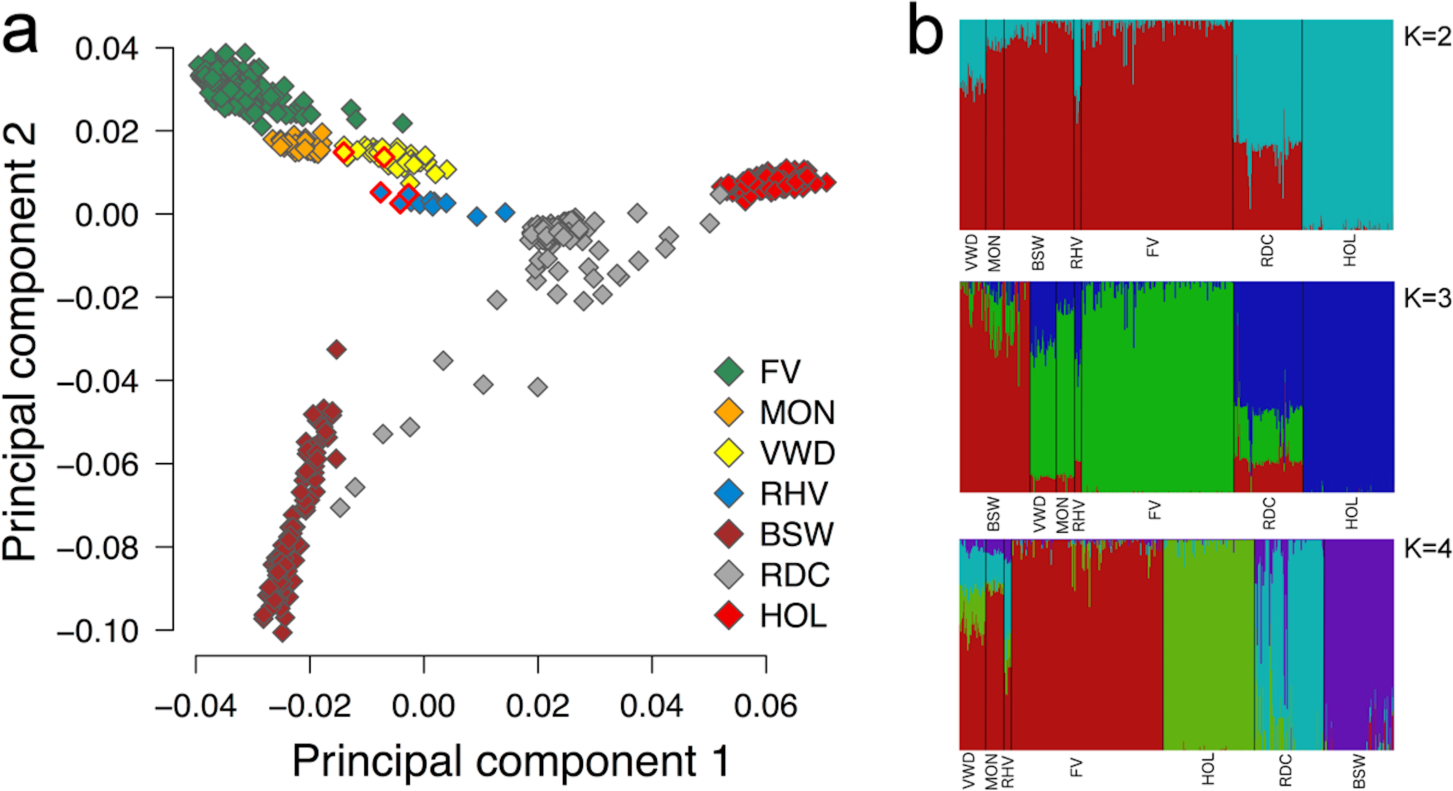
Genetic clustering of 714 animals from seven cattle breeds. Plot of the top two principal components of the genomic relationship matrix (a). Symbols with red frames represent five calves with epidermolysis bullosa from the Vorderwald (VWD) and Rotes Höhenvieh (RHV) cattle breed. b) Population structure of seven cattle populations obtained using the *fastSTRUCTURE* software tool. Different colours indicate ancestry proportions for 2, 3 and 4 predefined genetic clusters (b). FV: Fleckvieh, MON: Montbéliarde, VWD: Vorderwald, RHV: Rotes Höhenvieh, BSW: Brown Swiss, RDC: Red Dairy Cattle, HOL: Holstein Friesian.

The EB-associated haplotype was not present in 659 animals from breeds other than Vorderwald and Rotes Höhenvieh cattle. It turned out that Piras inherited the non-associated haplotype from its Montbéliarde sire while the EB-associated haplotype was transmitted from its mother that has no documented Montbéliarde ancestry.

## Discussion

Twenty-five calves with autosomal recessive epidermolysis bullosa (EB) were noticed in the local German Vorderwald cattle population. Genomic regions underlying recessive conditions are usually identified by comparing genome-wide marker data between affected and unaffected individuals (*e.g.*, [5]). We could not apply case-control association testing for the mapping of EB because only two affected animals were genotyped [28]. Since genetic material from additional cases had not been collected, we resorted to include genotypes of obligate mutation carriers, *i.e.*, parents of affected calves, in our association study. We additionally genotyped artificial insemination bulls that had no descendants with EB because they are less likely to carry the disease-associated haplotype. Using the presumed number of copies of the defective haplotype as response variable in a linear regression model allowed us to map EB to BTA22. Both diseased calves were identical by descent for a 21.26 Mb haplotype at the distal end of BTA22 corroborating that this region is associated with EB. Our association study was successful because we were able to unambiguously derive the haplotype state of unaffected individuals. The haplotype state of healthy animals may be less obvious for diseases with *e.g.*, an unknown mode of inheritance or genetic heterogeneity. However, our results evidence that haplotype-based association mapping may offer a useful tool for pinpointing genomic regions underlying Mendelian traits when a sufficient number of affected individuals is not immediately available [29, 30].

The analysis of whole-genome re-sequencing data of 1686 cattle revealed that a nonsense mutation (rs876174537) in the *COL7A1* gene segregates with EB. Sequence variants in *COL7A1* cause epidermolysis bullosa in various species [19, 31, 32, 33, 34]. Interestingly, Menoud et al. [19] detected the rs876174537 T-allele in three calves with dystrophic EB from the Rotes Höhenvieh cattle breed. The exchange of genetic material between Vorderwald and Rotes Höhenvieh cattle may explain the manifestation of the same recessive condition in two distinct cattle breeds [13, 16]. Deleterious alleles that segregate across populations may also originate from unknown ancient ancestors, predating the separation of the two breeds [35]. Since the diseased calves from both breeds are identical by descent for a large haplotype (8.72 Mb) encompassing the defective allele, a recent common ancestor is more likely. However, incomplete pedigree information from affected calves particularly from Rotes Höhenvieh cattle [19, 26], precluded the identification of common ancestors. Two sources of the EB-associated haplotype are plausible: first, a mutation carrier from the Vorderwald cattle breed may have been used for cross-breeding in Rotes Höhenvieh cattle or *vice versa*. However, to the best of our knowledge, an admixture of both breeds had not been documented. Second, the mutation may originate from an unknown foreign ancestor of both breeds. While Vorderwald cattle experienced substantial admixture with dairy and dual purpose breeds in the past 40 years [2], little is known regarding the genetic constitution of Rotes Höhenvieh cattle. Our population-genetic analyses revealed, unexpectedly, high genetic similarity between Rotes Höhenvieh, Vorderwald and Montbéliarde cattle indicating that common haplotypes may segregate across these breeds. However, a Montbéliarde origin of the rs876174537 T-allele is less likely, because the VorderwaldxMontbéliarde crossbred bull Piras inherited the EB-associated haplotype from its mother that has no Montbéliarde ancestry. A Vorderwald calf with clinical features similar to those observed in our study was reported in 1972 [36]. It is plausible that the nonsense mutation in *COL7A1* occurred several generations ago in an unadmixed Vorderwald animal before the introgression of foreign haplotypes began.

Between five and ten Vorderwald bulls are annually selected for artificial insemination. The identification of the mutation causing EB now allows for the reliable detection of animals carrying the defective allele. In 79 bulls that were used for artificial insemination in the past 20 years, among them seven sires of calves with EB, the frequency of the rs876174537 T-allele was 5%. However, the widespread use of an unnoticed mutation carrier for artificial insemination may result in a rapid increase of the frequency of the deleterious allele particularly when considering the small size of the Vorderwald cattle population [9, 11]. Since the defective allele was only found in the descendants of Piras, it seems advisable to refrain from using carrier animals for artificial insemination and select half-sibs without the defective allele instead. Applying this strategy should decrease the frequency of the defective allele in the Vorderald cattle population and prevent the birth of homozygous calves with a painful skin disease.

## Conclusions

A nonsense mutation in the *COL7A1* gene causes recessive epidermolysis bullosa in Vorderwald and Rotes Höhenvieh cattle. Since an admixture of both breeds had not been documented, the defective allele likely occurred several generations ago in an unknown common ancestor of Vorderwald and Rotes Höhenvieh cattle. Our findings demonstrate that deleterious alleles may segregate across cattle breeds without any documented admixture. The identification of the causal mutation now enables the reliable identification of carrier animals. Genome-based mating programs can avoid inadvertent matings of carrier animals thereby preventing the birth of homozygous calves suffering from a painful skin disease.

## Methods

### Animals

Twenty-five Vorderwald calves with EB were born between the years 2004 and 2015. The pedigree of the diseased animals was constructed using the *kinship2* R package [37]. The affected calves were descendants of nine sires and 24 dams. The number of progeny with EB ranged from 1 to 6 per sire. Breeding consultants collected ear tissue samples from two affected calves. In order to facilitate the mapping of the EB-associated genomic region, ear tissue and semen samples of dams (n=7) and sires (n=7), respectively, of another 23 diseased calves were collected by breeding consultants and employees of an artificial insemination centre. DNA from ear tissue and semen samples was prepared following standard DNA extraction protocols.

### Genotyping, quality control and haplotype inference

Two calves with EB and 14 parents of another 23 affected calves were genotyped with the Illumina BovineSNP50 v2 BeadChip comprising genotypes at 54,609 SNPs. Twenty-seven healthy artificial insemination bulls from the Vorderwald cattle breed were genotyped with the Illumina BovineSNP50 v1 BeadChip comprising genotypes at 54,001 SNPs. The genotype data were combined and 52,339 SNPs that were a common subset of both datasets were considered for further analyses. The per-individual call-rate ranged from 95.10% to 99.87% with an average call-rate of 98.59%. The chromosomal positions of the SNPs corresponded to the UMD3.1 assembly of the bovine genome [38]. We did not consider 412 SNPs with unknown chromosomal position for further analyses. Following quality control using *plink* v1.90 [39] (minor allele frequency above 0.5%, no deviation from the Hardy-Weinberg equilibrium (P>0.0001), per-SNP and per-individual call-rate higher than 95%), 43 animals (2 affected calves, 14 parents of affected calves and 27 control animals) and 41,436 SNPs were retained for association testing. The *findhap* software (version 3) [40] was used to impute sporadically missing genotypes and to infer haplotypes while accounting for pedigree information.

### Haplotype-based association testing

A sliding window consisting of 20 contiguous SNPs (corresponding to an average haplotype width of 1.23±0.39 Mb) was shifted along the chromosomes in steps of 5 SNPs. Within each sliding window (N=8207), all haplotypes (N=57,837) with a frequency above 5% were tested for association with EB using a linear regression model that included a correction term to account for the relationship among the animals: 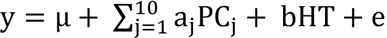, where y was a vector of phenotypes (coded as 2, 1 and 0 for calves with EB, obligate mutation carriers and control animals, respectively), µ was the intercept, PC_j_ were j=10 principal components of the genomic relationship matrix that was built based on 40,792 autosomal SNPs using the *GCTA* software tool [41], a and b were regression coefficients for ten principal components and the haplotype (HT) tested, respectively, where HT was coded 0, 1 and 2 for non-carrier, heterozygous and homozygous animals, respectively, and e was a vector of residuals that were assumed to be normally distributed. Association testing was implemented in R (see Additional file 2) and haplotypes with P values less than 8.6x10^−7^ (Bonferroni-corrected significance threshold for 57,837 independent tests) were considered as significantly associated with EB.

### Generation of sequence data

Genomic DNA was prepared from ear tissue and semen samples of an affected calf, its parents and another heterozygous haplotype carrier following standard DNA extraction protocols. Paired-end libraries were prepared using the paired-end TruSeq DNA sample preparation kit (Illumina) and sequenced using the HiSeq 2500 instrument (Illumina). The resulting reads were aligned to the University of Maryland reference sequence of the bovine genome (UMD3.1 [38]) using the *BWA-MEM* algorithm [42, 43]. Individual files in SAM format were converted into BAM format using *SAMtools* [44]. Duplicate reads were marked with the MarkDuplicates command of *Picard Tools* [45].

### Variant calling, imputation and identification of candidate causal variants

Single nucleotide and short insertion and deletion polymorphisms were genotyped in four animals simultaneously using the multi-sample approach implemented in *mpileup* of *SAMtools* along with *BCFtools* [44]. *Beagle* phasing and imputation [46] was used to improve the primary genotype calling by *SAMtools*. To identify mutations compatible with recessive inheritance, all polymorphic sites that were detected within the 21.26 Mb segment of extended homozygosity were filtered for variants that were homozygous for the non-reference allele in the affected calf and heterozygous in three haplotype carriers. The genotypes of thus identified candidate causal variants were validated in 1682 animals from various bovine breeds that were available from run 5 of the 1000 bull genomes project [47] and our in-house bovine genome database [35], respectively. Sequence variants that were homozygous for the non-reference allele in healthy animals were excluded as plausible candidate causal variants. The functional consequences of sequence variants compatible with recessive inheritance were predicted using the *Variant Effect Predictor* tool from Ensembl [48, 49].

### Validation of candidate causal variants

We obtained genotypes for the rs876174537 polymorphism in 88 animals including 2 affected calves, 14 parents of affected calves and 72 healthy artificial insemination bulls that were born between 1982 and 2014. DNA of artificial insemination bulls was prepared from semen straws following standard DNA extraction protocols. PCR primers (GGCTGATCGTCTTTGTCACC (forward) and TCAGTCCTGATCCCCAACTC (reverse) as detailed in [19]) were designed to obtain genotypes by Sanger sequencing. Genomic PCR products were sequenced using the BigDye® Terminator v1.1 Cycle Sequencing Kit (Life Technologies) on the ABI 3130×1 Genetic Analyzer (Life Technologies).

### Population genetic analyses

We merged the genotype data of 43 Vorderwald cattle with genotype data of 671 animals from another six cattle breeds (Fleckvieh, Holstein, Rotes Höhenvieh, Montbéliarde, Red Dairy Cattle, Brown Swiss) to perform population genetic analyses. Array-derived genotypes of 250 and 150 randomly selected Fleckvieh and Holstein animals, respectively, were available from previous studies [9, 50]. The genotypes of 12 cattle from Rotes Höhenvieh including three calves with dystrophic EB were obtained from Menoud et al. [19]. Genotypes of 115, 114 and 30 animals of the Brown Swiss, Red Dairy and Montbéliarde cattle breeds were extracted from whole-genome sequence variants provided by the 1000 bull genomes project [47]. The final data set consisted of 714 animals and 37,608 autosomal SNPs that were a common subset of all sources. The genomic relationship matrix among the animals was built and principal components were obtained using the *GCTA* software tool [41]. The estimation of ancestry proportions was performed with the *fastSTRUCTURE* software tool [51] using between two and seven predefined populations (K). In addition, haplotypes consisting of 60 contiguous SNPs from the Bovine SNP50 Bead chip surrounding the rs876174537 polymorphism were available for 576 Montbéliarde relatives of the bull Piras.

## Abbreviations

BTA: Bos taurus; EB: epidermolysis bullosa; N_e_: effective population size; RHV: Rotes Höhenvieh; SNP: single nucleotide polymorphism; VWD: Vorderwald

## Declarations

### Acknowledgements

We thank Dr. Franz Maus from Landratsamt Schwarzwald-Baar-Kreis for providing ear tissue samples of diseased calves and parents of diseased calves. We acknowledge Rinderunion Baden-Württemberg e.V. for providing semen samples of artificial insemination bulls. We thank Aurélian Capitan for providing haplotype data of Montbéliarde cattle and the 1000 Bull Genomes consortium for providing whole-genome sequence variants for key ancestors of various bovine breeds.

### Availability of supporting data

All relevant data are included within the article and its additional files.

### Authors’ contributions

SA and CW performed molecular-genetic analyses; CW and RF performed next-generation sequencing and analyzed sequencing data; HH provided genotype and pedigree data of VWD cattle; CD and JT provided genotype data of RHV cattle; HP performed the statistical analyses, wrote the manuscript and conceived the study. All authors read and approved the final manuscript.

### Ethics approval and consent to participate

Two calves with EB were inspected by accredited veterinarians. Due to the severe disease manifestations with no prospect of improvement, both animals were euthanized and ear tissue samples were collected for genetic investigations. The decision to euthanize the calves was solely at the discretion of the veterinarians. None of the authors of the present study was involved in the decision to euthanize the calves. No ethical approval was required for this study.

## Additional Files

### Additional File 1

File format:.txt

**Title**: Annotation of variants compatible with recessive inheritance

Description: The functional consequences of 702 sequence variants compatible with recessive inheritance of epidermolysis bullosa were obtained from Ensembl using the *Variant Effect Predictor* tool.

### Additional File 2

File format:.tar.gz

**Title**: R script implementing the GWAS algorithm applied to map EB

Description: R script (haplo_GWAS.r) and an example dataset consisting of haplotypes for chromosome 22 (phased_BTA22), the positions of the SNPs located on BTA22 (BTA22.map) ten principal components (vwd.eigenvec) and phenotypes (pheno_epiderm_bullosa) for two affected calves, 14 parents and 27 control animals. The script was tested in an unix environment.

